# Bulk-level maps of pioneer factor binding dynamics during Drosophila maternal-to-zygotic transition

**DOI:** 10.1101/2024.09.06.611749

**Authors:** Sadia Siddika Dima, Gregory T. Reeves

## Abstract

Gene regulation by transcription factors (TFs) binding cognate sequences is of paramount importance. For example, the TFs Zelda (Zld) and GAGA factor (GAF) are widely acknowledged for pioneering gene activation during zygotic genome activation (ZGA) in *Drosophila*. However, quantitative dose/response relationships between bulk TF concentration and DNA binding, an event tied to transcriptional activity, remain elusive. Here, we map these relationships during ZGA: a crucial step in metazoan development. To map the dose/response relationship between nuclear concentration and DNA binding, we performed raster image correlation spectroscopy, a method that can measure biophysical parameters of fluorescent molecules. We found that, although Zld concentration increases during nuclear cycles (ncs) 10 to 14, its binding in the transcriptionally active regions decreases, consistent with its function as an activator for early genes. In contrast, GAF-DNA binding is nearly linear with its concentration, which sharply increases during the major wave, implicating it in the major wave. This study provides key insights into the properties of the two factors and puts forward a quantitative approach that can be used for other TFs to study transcriptional regulation.

## Introduction

In *Drosophila*, ZGA begins with the transcription of a handful of genes during the minor wave of ZGA, followed by the major wave when thousands of genes are transcribed^1–4^. The TF Zld has the ability to bind nucleosomal DNA^5,6^ and subsequently to facilitate the binding of other TFs^7–13^: the two defining features of a special class of TFs known as pioneer factors^14–16^. The maternally encoded TF GAF also possesses pioneer-like properties^10,11,17–20^.

The role of TFs binding DNA for gene regulation is widely accepted in the context of development and homeostasis^21–23^. However, the relationship between the concentration of TFs such as these pioneer factors and gene expression remains an open question^24,25^. Fluorescent imaging of live and fixed tissues can give relative concentrations of TFs, which are often used to predict TF activity by assuming some nonlinear relationship between concentration and activity^26–32^. However, validation of these relationships using dynamic, quantitative data in live cells is currently lacking. Furthermore, the binding of the TFs to DNA –required for transcriptional regulation – is dependent not only on their nuclear concentration, but also on factors such as chromatin accessibility and saturation kinetics. On the other hand, recent studies have identified hubs of TF binding at DNA sites, which may be more direct measures of TF activity^33–36^. However, the relationship between these hubs and the bulk TF concentration (experimentally accessible from fluorescence measurements) is unknown. Thus, knowing only total concentration is inadequate to predict TF binding, and direct measurements of TF hubs lack generalizability; an input/output map between the two, based on quantitative measurements of concentration and binding, is required.

To bridge this gap, we used raster image correlation spectroscopy (RICS) to quantify the concentration and binding of the pioneer-like factors Zld and GAF in live embryos over the course of nc 10-14^37–42^. These measurements allowed us to construct dose/response relationships which suggest that the binding of Zld to transcriptionally active sites is dependent on factors other than its concentrations alone, whereas the GAF concentration is the primary driver of its binding. Furthermore, we found that GAF must bind and saturate its sites in the inactive regions of the DNA before it can bind to active DNA sites, resulting in a delay in its role in ZGA. Similar approaches can be used to obtain comprehensive quantitative picture of dynamics of other TFs for gene regulation studies.

## Results and discussion

Raster-scanned confocal images have a fast-scanning direction (pixel-to-pixel) and a slow-scanning direction (line retracing; Figure 1A). RICS analysis uses intensity fluctuations of GFP-tagged molecules, correlated in time and space, to build an autocorrelation function (ACF) in two dimensions, corresponding to the fast (Δx) and slow (Δy) scanning directions, respectively (Figure 1B; see Methods). The amplitude of the ACF, *A*, is inversely proportional to the GFP concentration (Figure 1C), while the shape of the ACF, especially in the slow direction, is determined by the fraction of GFP that is freely diffusible vs. that which is immobile (Figure 1D). Furthermore, cross-correlations in the fluctuations between GFP and RFP (tagged to His2Av) allows us to calculate the fraction of GFP-tagged molecules that are bound to the same structure as the His2Av-RFP (likely to be DNA^43,44^; Figure 1E). Therefore, RICS allows us to quantify not only the absolute concentration of GFP molecules, but also the fraction freely diffusible, the fraction immobile, and the fraction correlated to His2Av (Figure 1F).

**Figure 1.**
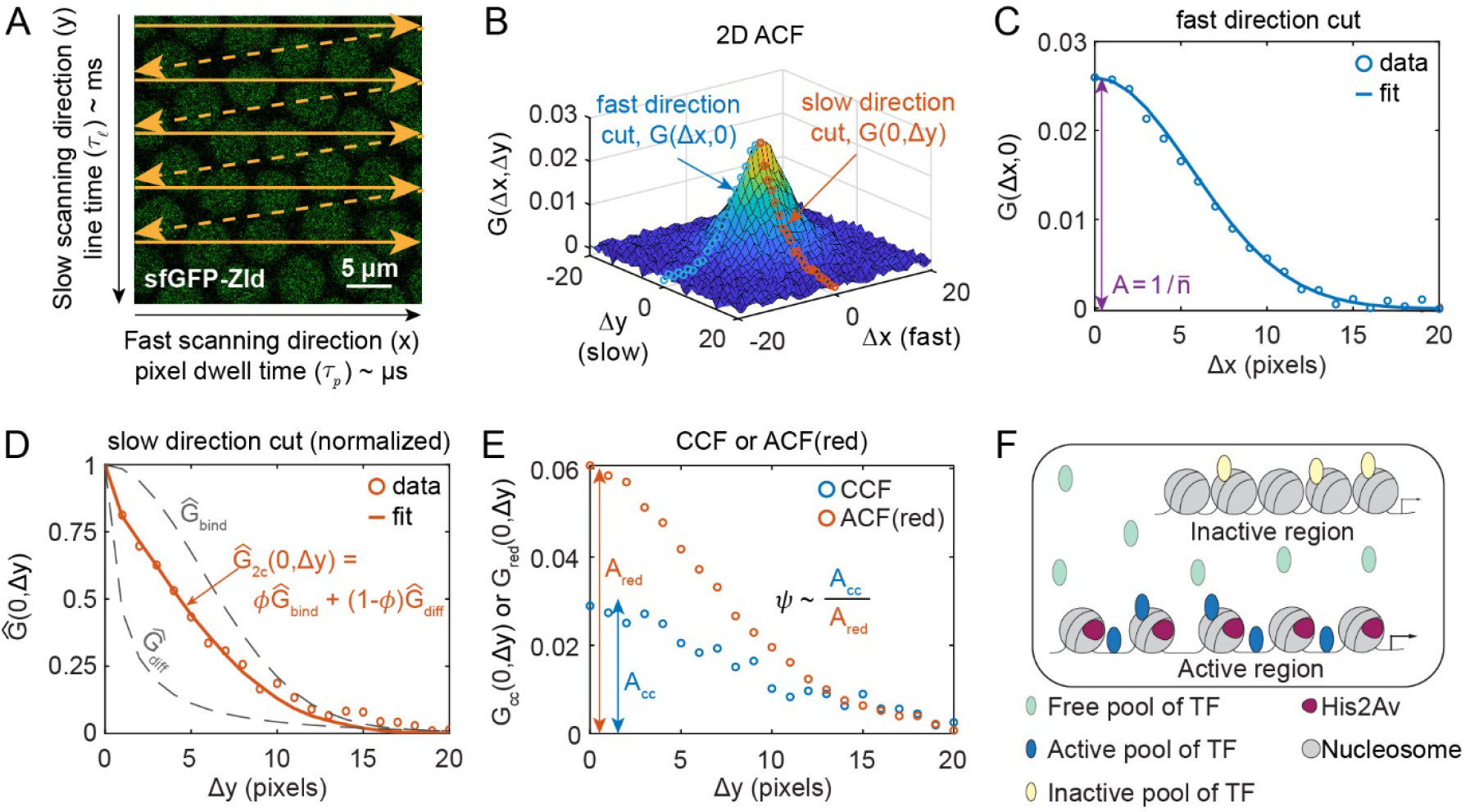
Raster Image Correlation Spectroscopy (RICS). (A) Laser scanning confocal microscopes build images by raster scan, with a fast scanning direction (x direction, solid arrows), and a slow scanning direction (y direction) due to line retracing (dotted arrows). The time between horizontally-neighboring pixels is ∼microseconds, while that between vertically-neighboring pixels is ∼milliseconds. Image: mid-nc14 embryo expressing sfGFP-Zld. (B) Two-dimensional ACF from embryo depicted in (A). Cuts along the fast (blue circles) and slow (orange circles) directions are depicted. (C) Cut of ACF along the fast direction. Solid curve: fit of Gaussian-shaped PSF, used to estimate the ACF amplitude, *A*. (D) Plot of the slow direction data (circles) and the fit to the slow direction (solid curve), composed of a linear combination between two ACFs (gray dotted curves): an immobile ACF 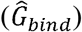and a diffusible ACF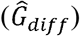. The linear combination weight is *ϕ*, the fraction immobile. (E) Plots of cuts of the cross correlation function (blue circles) and of the ACF in the red channel (orange circles). The ratio of the amplitudes (*A*_*cc*_*/A*_*red*_) is proportional to *ψ*, the fraction bound in active regions of the DNA. (F) Illustration of the different pools of TF: freely diffusible (light blue), bound to active regions (dark blue), and bound to inactive regions (yellow). His2Av (red) is associated with the active regions of DNA.

### Zld levels bound to DNA decrease while nuclear concentration increases

We performed RICS analysis on the nuclei of blastoderm-stage embryos expressing sfGFP-Zld^45^ (Figure 2A and Video S1) to measure the dynamics of the nuclear concentration of Zld over time. We found the total nuclear concentration of Zld increased from one nuclear cycle to the next during nc 10 to 14, suggesting an increase in the Zld levels with time (Figure 2B). Increase in Zld levels have been reported previously using immunoblotting^46,47^. Furthermore, the nuclear concentrations of Zld vary significantly within each nc, as more sfGFP-Zld enters nuclei after mitosis and fills it during the interphase. The longer duration of nc 14 allows Zld levels to reach a steady state, unlike the other, shorter ncs. In contrast, we observed that the nuclear Zld concentration drastically decreases from the end of one interphase to the beginning of the next (during mitosis), in agreement with previous observations that, unlike most pioneer factors, Zld is not mitotically retained on the chromosomes^36^.

**Figure 2.**
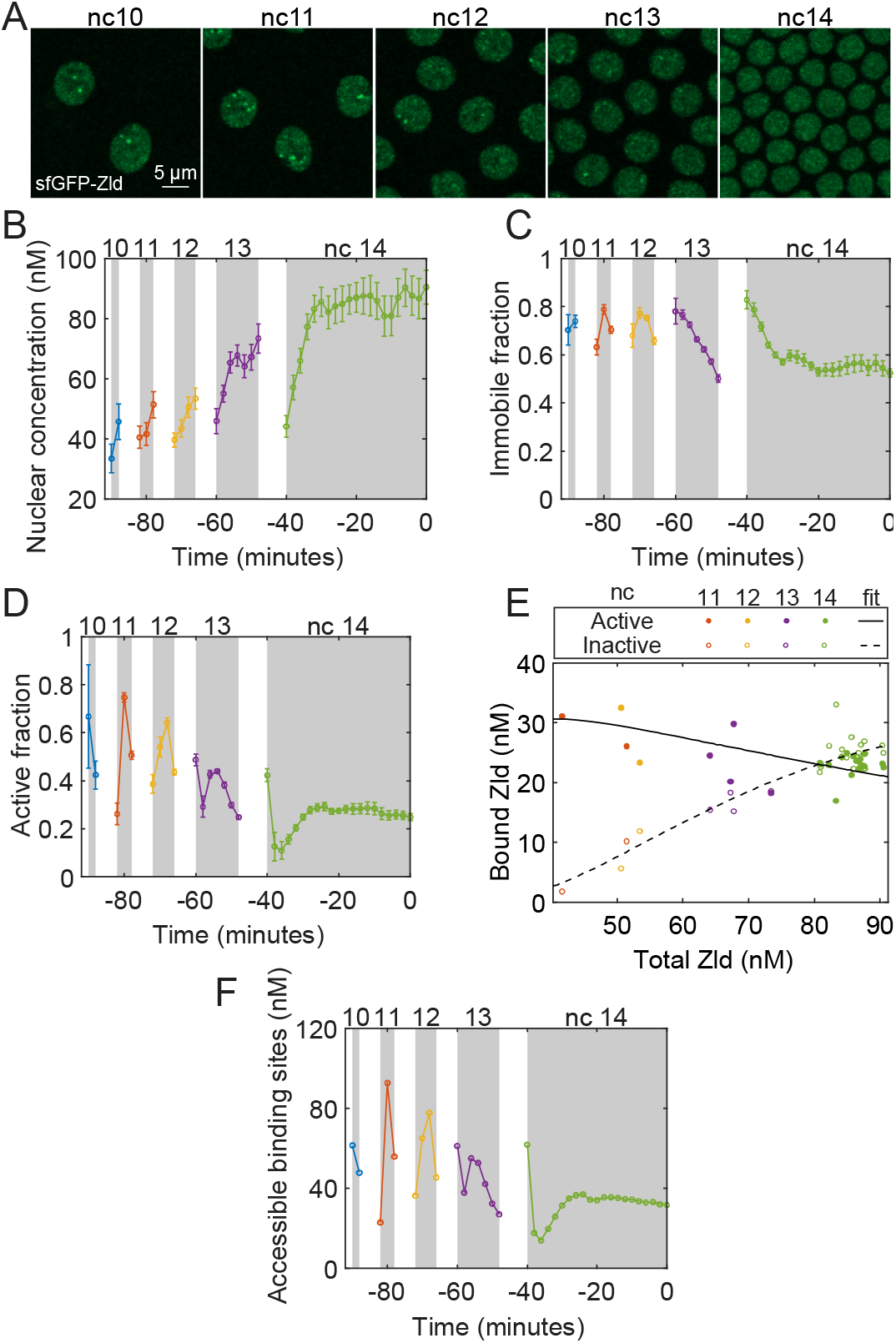
Quantification of the biophysical parameters and dynamics of Zld. (A) Representative images of sfGFP-Zld embryo used for RICS analysis from nc 10 to nc 14 as marked. (B-D) Dynamics of different pools of sfGFP-Zld from nc 10 until gastrulation including total nuclear concentration (B), immobile fraction (C), and active fraction (D). Curves: mean values (n = 20 embryos). Error bars: S.E.M. (E) Dose/response map between total nuclear concentration of Zld and the immobile (both active and inactive) concentration of Zld. (F) Change of Zld sites accessible for binding over time. See also Figure S1 and Video S1.

RICS analysis also allowed us to determine *ϕ*, the fraction of Zld which is immobile (or nearly so) due to either binding to immobile structures (such as DNA) or forming large aggregates having very low diffusivity (Equation 4). The immobile fraction of Zld remains nearly the same from nc 10-13, after which it reaches a steady state at a lower value of ∼0.5 at nc 14 (Figure 2C). Fifty percent of the Zld population was found to be immobile in single-molecule imaging^34^.

In each nc, at the beginning of interphase, the immobile concentration of Zld, calculated as the product of total nuclear concentration and the immobile fraction, increases perhaps because the chromatin decondenses and Zld binds the replicated DNA^5,18,48,49^ and it starts decreasing after reaching a maximum as chromatin starts condensing before mitosis (Figure S1A). This is consistent with the rapid formation of dynamic hubs of Zld after mitosis^34^, the hubs likely being a population included in our immobile concentration measurements. It should be noted that the presence of hubs results in Zld particles with higher brightness than Zld monomer, given Zld is unlikely to form multimers^45,46,50,51^. This in principle could alter our measurements of total Zld nuclear concentration and of the immobile fraction. We have quantified the average particle brightness and found it to be roughly constant over time, suggesting that hubs of Zld have little effect on our measurements (see Methods and Figure S2).

We also used cross-correlation RICS (ccRICS) to determine *ψ*, the fraction of the pioneer factors that correlate with His2Av-RFP (Equation 9 in Methods). In *Drosophila*, the enrichment of the histone variant His2Av has been found to correlate with transcriptional potential^52,53^, and therefore, the pool correlated with His2Av-RFP (hereafter referred to as the active pool) is likely responsible for pioneering gene activation. As the immobile fraction always measured as larger than the active fraction (Figures 2C, D), we inferred that the immobile fraction is composed of two pools: one that correlates with His2Av-RFP and one that does not. Hereafter, the pool that does not correlate with His2Av but is immobile, either due to binding to inactive chromatin regions or due to low diffusivity from the formation of large clusters, is referred to as the inactive pool. We found the active fraction decreases from nuclear cycle to nuclear cycle (Figure 2D). Consequently, the active concentration of Zld, which is the product of the total nuclear concentration and the active fraction of Zld, also decreases, albeit weakly due to the increasing total nuclear concentration (Figure S1B). Our results agree with the slight reduction observed in the number if Zld peaks from nc 13 to 14 in ChIP-Seq^54^. Roughly 5 min into nc 14, the concentration of active Zld reaches a steady value around 20-25 nM (Figure S1B).

Our measurements lend themselves to a general analysis in which the relationship between Zld total nuclear concentration and its binding could be constructed. According to standard thermodynamic equilibrium binding models (see Methods), an increase in the total Zld concentration should lead to an increase in the bound Zld concentration. However, our data suggest that the active concentration of Zld is a weakly *decreasing* function of total Zld (Figure 2E) in which nc 10-12 has a high active concentration of Zld despite the low total Zld concentration, while in nc 14, there is a high total Zld concentration and a slightly lower active concentration. As such, our results suggest that Zld binding is not influenced by Zld concentration alone, and additional factors, such as chromatin structure, must be taken into account. Using our data and ChIP-seq results reported by Harrison et al.^54^, the dissociation constant (Kd) for Zld can be estimated to be ∼17 nM (see Methods). Using this value, fitting a Hill function to our data suggests that the Zld sites accessible for binding reduce over time (Figure 2F), which might result from the chromatin becoming more densely packed during the later cycles^8,54,55^. On the other hand, the inactive concentration of Zld increases with time (Figure S1C), and thereby is correlated with the increase in total Zld concentration as well, and is functionally consistent with standard thermodynamic equilibrium binding models, suggesting the inactive fraction may result from binding that is saturating in nature. Fitting a Hill function with a Hill coefficient of 2 yielded the Kd of ∼24 nM and a max concentration of 34 nM (Figure 2E). These values of Kd suggest that the binding of Zld to DNA or to immobile structures/aggregates is moderately strong. Overall, the dose/response relationship between total and bound Zld concentration is consistent with the previously-observed hubs of Zld bound to DNA. Zld molecules within these hubs have short residence times on DNA (*τ ∼* 5 *s*^34^). Although this is close to our estimation of the residence time (see methods), the slight reduction observed in the case of hubs might be due to the high local concentration of Zld within the hub^34^, or the local Zld diffusivity^56^.

Our data suggest that Zld may not retain its ability to pioneer chromatin accessibility throughout ZGA. Consistent with this, it was recently shown that, in larval type II neuroblasts, Zld binding is influenced by chromatin accessibility^57^. Thus, our observed decrease in the active concentration may imply that Zld primarily regulates transcription during the minor wave. Indeed, the removal of Zld activity affects early gene expression patterns, resulting in a delay in, but not complete loss of the transcription of patterning genes^47,58^. For example, in the Dorsal (Dl) network, the target genes *sna, twi, rho, brk, sog* are delayed in embryos lacking maternal Zld^47,58^, and the removal of Zld binding sites from the *sog* neurogenic ectoderm enhancer results in delayed and sporadic *sog* expression^9,59^. Furthermore, mathematical models that account for decreasing Zld levels concurrent with increasing Dl levels ^60–62^ can explain Dl target gene expression^27,63^. Therefore, our results suggest that, during the minor wave of ZGA (nc 10-13), a high concentration of bound Zld might be needed to facilitate the binding of low Dl levels. On the other hand, during the major wave at nc 14, either the high level of Dl, or the presence of other pioneer-like factors, may be sufficient to drive Dl binding and target gene expression when the bound Zld levels have decreased. This raises the question of whether other factors are present during the major wave that may continue to facilitate the binding of other developmental TFs.

### GAF levels increase suddenly in nc 14

GAF, encoded by the *Trithorax-like* (*Trl*) gene, plays an essential role in ZGA along with Zld^17,64^. GAF motifs along with Zld motifs are enriched in the highly occupied target (HOT) regions characterized by open chromatin and bound by many TFs^65–67^. GAF binds to clustered motifs^68–70^ and possesses the properties of a pioneer factor, such as the ability to bind nucleosomal DNA and to create regions of chromatin accessibility by functioning with chromatin remodelers to facilitate the binding of other TFs^17,20,71–74^.

To measure the dynamics of the nuclear concentration of GAF over time, we performed RICS analysis on the nuclei of blastoderm-stage embryos expressing GAF-sfGFP^17^ (Figure 3A and Video S2). The results suggest that the total nuclear concentration of GAF remained nearly constant and very low (∼5-10 nM) from nc 10-13, then showed a sudden increase during nc 14 to ∼30 nM (Figure 3B). We saw that the nuclear concentration at the beginning of an interphase was similar to that at the end of the previous interphase (Figure 3B). This observation is consistent with previous work showing GAF is mitotically retained on the chromosomes^17^, and is in contrast to Zld (Figure 2B), which is not mitotically retained^36^.

**Figure 3.**
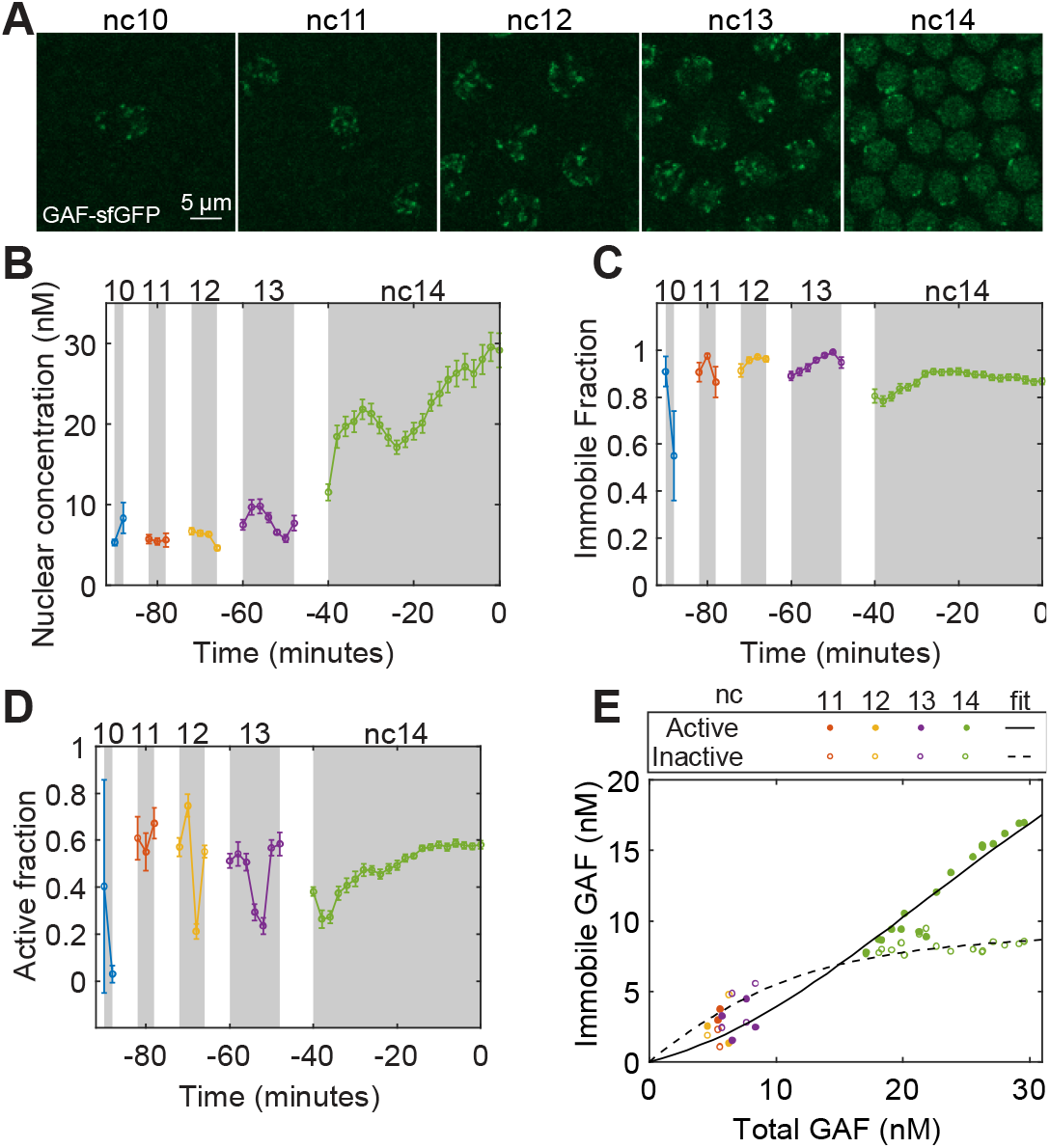
Quantification of the biophysical parameters and dynamics of GAF. (A) Representative images of GAF-sfGFP(C) embryo used for RICS analysis from nc 10 to nc 14 as marked. (B-D) Dynamics of the parameters from nc 10 until gastrulation including total nuclear concentration (B), immobile fraction (C), and active fraction (D). Curves: mean values (n = 19 embryos). Error bars: S.E.M. (E) Dose/response map between total nuclear concentration of GAF and the immobile (both active and inactive) concentration of GAF. See also Figure S3 and Video S2.

The immobile fraction of GAF remained nearly the same during nc 10-13, then decreased slightly during nc 14 (Figure 3C). The sudden increase in GAF total nuclear concentration, together with only a slight decrease in its immobile fraction, resulted in an increase in the immobile concentration of GAF during nc 14 (Figure S3A). Although the active fraction of GAF, the pool expected to be bound near the actively transcribed genes, remained approximately the same on average from nc 10-14, it varied significantly within each nc (Figure 3D).

As with Zld, we sought to use our data to map the relationship between GAF nuclear concentration and binding. We noted that, unlike Zld, both active and inactive concentrations of GAF increase with an increase in the total concentration of GAF (Figure 3E). Our results agree with the increase in GAF peaks from nc 9 to 14 observed previously^17^. Both pools appeared to be saturating functions of total GAF concentration, with the inactive pool saturating quickly at a low overall concentration (Figure 3E and Figures S3B-C). In contrast, the active pool required higher levels of total GAF to saturate and had a higher capacity. Fitting Hill functions to the data bore out these observations: the inactive pool had a Kd of 0.4 nM and a max concentration of 9 nM, while the active pool had a Kd of 5 nM and a max concentration of 35 nM. The Kd values represent a strong affinity of GAF for the binding sites, roughly 4-fold and 60-fold greater than for Zld, consistent with previous reports of stable GAF/DNA binding^72,75,76^. Interestingly, the higher affinity/lower capacity inactive pool may in part explain the sudden increase in the lower affinity/higher capacity active pool in nc 14. It appears that, as GAF concentration slowly increases, the majority is apportioned to the inactive pool due to its high affinity. However, because of the low capacity of the inactive pool, upon entering nc 14, the active pool is suddenly able to increase. If the active pool corresponds to transcriptionally active regions of the DNA (*i*.*e*., euchromatin), the relative affinities and capacities ensure that GAF acts as a pioneer factor solely during the later, major wave of ZGA. Furthermore, it has been shown that GAF associates to the GA/CT-rich repeats in the heterochromatin regions throughout cell cycles^70,77^, potentially driving transcriptional silencing and the euchromatin association drives activation during ZGA^70^. As such, the inactive pool may represent GAF bound to heterochromatin. As with Zld, GAF hubs with higher brightness could affect our measurements. Quantification of the brightness suggested small corrections to our data, which did not affect our overall conclusions (see Methods and Figure S4).

Our measurements of the biophysical parameters of the pioneer factors and their dynamics allowed the construction of input/output maps between total concentration of the factors and their DNA binding. While a subset of the pioneer factor bound regions has been found to stay inaccessible^78,79^, binding of pioneer factors to DNA is an indispensable step for establishing and maintaining chromatin accessibility. Following this, binding of a correct combination of TFs can lead to the activation of the accessible regions^80^. Using the input/output maps we can infer that Zld acts as a global activator of early genes enabled by the high active concentration of Zld during the minor wave of ZGA, and GAF comes into play during the major wave, as indicated by the high active concentration of GAF during this time (Figure 4). This is supported by the fact that the regions that gain accessibility early during ZGA are enriched for Zld binding, whereas the ones that gain accessibility late are enriched for GAF binding^18^. A similar quantitative approach, in which accurate measurements of biophysical parameters of TFs and their dynamics enable the construction of input/output map between TF concentration and transcriptional activity, is expected to aid other gene expression studies.

**Figure 4.**
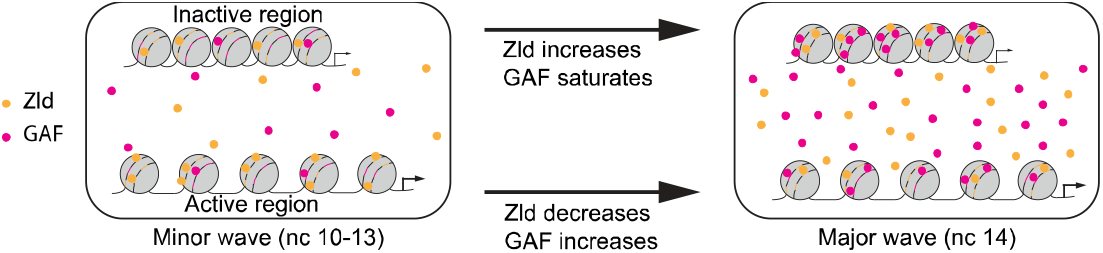
Role of pioneer factors Zld and GAF during *Drosophila* ZGA. During the minor wave of ZGA high concentration of Zld (yellow circles) bound to its sites (yellow lines) in the active regions allows it to act as the main activator of early genes. This active bound concentration reduces during the major wave when the concentration of GAF (magenta circles) bound to the sites (magenta lines) in the active regions increases, after saturating the sites in the inactive regions, allowing GAF to act as the main activator of the genes that begin expressing during the major wave.

## Acknowledgements

We thank Melissa M. Harrison for kindly providing fly stocks and for helpful discussion of the manuscript. This work was supported in part by NSF grant MCB-2105619.

## Author contributions

S.S.D and G.T.R designed the experiments and wrote the manuscript. S.S.D conducted the experiments, made Zld and GAF mathematical models. G.T.R developed the RICS analysis pipeline.

## Data and resource availability

Image analysis pipeline: https://github.com/gtreeves/RICS_timecourse_pipeline All the image files (.czi) will be deposited in the “Texas Data Repository”.

## Methods

### *Drosophila* strains

The fly strain used for the Zld imaging is *sfGFP-zld; His2Av-RFP (II)*. The *sfGFP-zld* mutant allele was generated using Cas9-mediated genome engineering by Hamm et al.^45^ The fly strain used for the GAF imaging is *His2Av-RFP (II); GAF-sfGFP(C) (III)*. The *GAF-sfGFP(C) (III)* mutant allele was generated using Cas9-mediated genome engineering by Gaskill, Gibson et al.^17^

### Sample preparation

Flies were raised on standard cornmeal-molasses-yeast medium at 25°C. Fly cages were prepared with desired fly strains and kept at room temperature for 2 days before imaging. Grape juice agar plates streaked with yeast paste were placed onto the bottoms of the cages for oviposition. For imaging, the flies were allowed to lay eggs for 1 hour after which the plates were removed for embryo collection. The embryos were transferred from the plates to mesh baskets, dechorionated using bleach for 30 s, and washed with deionized water to remove residual bleach^81^.

### Live imaging

After dechorionation, the embryos were mounted in 1% low melting point agarose (IBI Scientific, IB70051) in deionized water on a glass bottom Petri dish (MatTek, P35G-1.5-20-C) and deionized water was poured into the Petri dish over the solidified low melting point agarose to cover the samples. The low melting point agarose rendered mechanical stability to the embryos while remaining transparent submerged in water^82,83^. All images were collected on an LSM 900 (Carl Zeiss, Germany) confocal laser scanning microscope. For the image acquisitions, C-Apochromat 40x/1.2 water immersion Korr objective, 488 nm laser for sfGFP, 561 nm laser for RFP, and GaAsP-PMT detector were used. The detector was operated at 650 V with 1x gain and 0% offset. Emission was detected in the range of 410-546 nm for sfGFP and 595-700 nm for RFP. All the images had 1024 × 1024 pixels resolution. The images were collected at 5x zoom, resulting in pixel size of 31.95 nm, and a pixel dwell time of 2.06 µs. This corresponds to a frame time of 5.06 s and a line time of ∼ 5 ms (ratio of frame time and number of rows in the image). Image acquisition was started when the embryos were at nc 10 and continued till gastrulation as indicated by the nuclear morphology.

### Raster image correlation spectroscopy (RICS) image analysis

RICS analysis, a derivative of fluorescence correlation spectroscopy, entails constructing autocorrelation functions (ACFs) from imaging data and fitting ACF models to these data-derived ACFs. We performed these analyses according to previous protocols^40,42^. In brief, live imaging time courses were background subtracted and divided into groups of 7-12 frames. The frames within a group were averaged together to create an averaged frame for that group, which was used for two purposes. First, the nuclei in the averaged frame were segmented according to a watershed algorithm. This segmented nuclear mask was used for each frame in the group. Second, the immobile variation within each frame was then removed by subtracting by the averaged frame on a pixel-by-pixel basis, then the scalar average intensity value of the averaged frame was added back^37,38^. Two dimensional (2D) RICS autocorrelation functions of the nuclear fraction of each frame were built using a fast Fourier transform protocol in Matlab, and these ACFs were averaged together for all frames in a given group. The result was a time series of 2D ACFs, each of which corresponded to a given grouping of 7-12 frames. Background subtraction was performed on the fly by examining the histogram of intensities and fitting a Gaussian to the lowest intensity pixels in the image.

### Experimental determination of axial displacement

We mounted diffraction-limited TetraSpeck beads (0.1 µm, T7279, Invitrogen) in 1% low melting point agarose on a glass bottom Petri dish following the same protocol as the embryos. Similarly, the agarose was immersed in deionized water upon solidification. The microscope parameters, such as the pixel size and frame time were identical as the RICS acquisitions. We acquired z-stacks with 0.05 µm distance between the slices. We acquired 6 z-stacks: 3 different locations in the agarose and 2 different batches of beads mounted in the agarose.

The beads were then detected in the average image of the z-stack. A 3D Gaussian was fitted to the intensity of the beads using fmincon. The centroid coordinates of each detected bead and the highest intensity z-plane for the corresponding bead were used as the initial guesses for the centers in x, y, and z directions. This was done separately for the two channels (Figures S5A-B). The difference between the centers (x, y, z) for the two channels as determined from the fit were calculated for each bead (Figures S5C-D). The full width at half maximum (FWHM) in the z-direction was calculated separately for the two channels (Figure S5E).

### Fitting RICS autocorrelation functions (ACFs) to estimate concentration and mobility

RICS autocorrelation functions generally contain two orthogonal pieces of information: the amplitude determines the concentration of the species and the shape determines the mobility. For a purely diffusing species, the theoretical, normalized ACF, 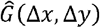, is the following:

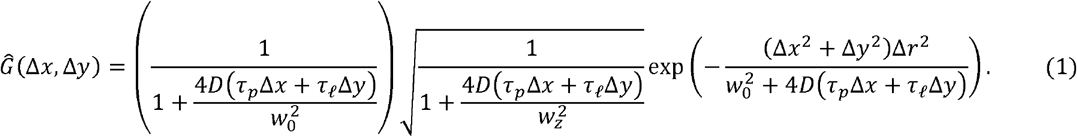

where *D* is the diffusivity. This equation has five microscope parameters: *w*_0_ and *w*_*z*_ are the radii of the point spread function (PSF) in the *xy* plane and the axial (*z*) direction, respectively; Δ*r* is the *xy* size of a pixel; and *τ*_*p*_ and *τ*_*l*_ are the pixel dwell time (determined by the scan speed) and line time (determined by a combination of the scan speed and number of pixels in the width of the image), respectively. The two independent variables, Δ*x*; and Δ*y*, are the pixel shifts in the fast and slow directions, respectively.

The non-normalized ACF includes the amplitude *A*, which is equal to the reciprocal of 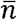 (the average number of molecules in the confocal volume):

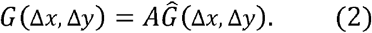

In practice, 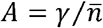, where 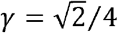 is a factor that accounts for the uneven illumination airy unit^41^. Note that 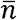 can be converted to 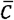, the average concentration in the confocal volume, *V*_*PSF*_, by 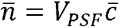, where 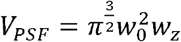.

The 2D ACFs for sfGFP-Zld or GAF-sfGFP for each time point were then used to fit two different models. First, the fast direction cut of the 2D ACF, *G*_*s*_(Δ*x*,0), was used to fit a Gaussian equation that approximates the microscope’s PSF. Because Δ*y* = 0 along this cut, and 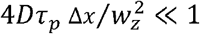, Equations (1-2) simplify to:

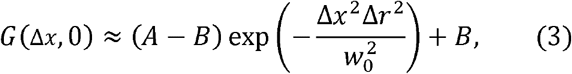

where, for robustness of fit, a small, adjustable background constant, *B*, was added and *w*_0_ was allowed to vary slightly. To avoid problems with background, *B* was constrained to have a magnitude less than 10^−3^, and in practice, it never exceeded 1% of the value of the amplitude, *A*. This first fitting step resulted in accurate estimates of the ACF amplitude, *A*.

Second, holding *A* fixed, the entire 2D ACF was then used to fit a two component model, *G*_2*c*_(Δ*x*, Δ*y*), which is a linear combination of a freely diffusing fraction and an immobile fraction^42^:

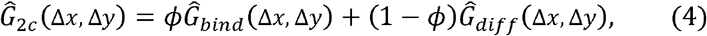

where 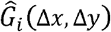 is given by Equation 1 (with 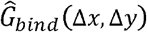 having *D* = 0 and 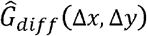 having *D* non-zero; see Figure 1D), the linear combination weight, *ϕ*, is the fraction immobile, and 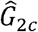 is the normalized two-component ACF, such that:

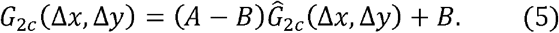

Note that the parameter *B* in Equation 5 may have a different value from the one found in Equation 3. The 2D ACF for His2Av-RFP for each time point was used to fit only the 2D PSF to get a measure of the ACF amplitude.

### Fitting ccRICS cross-correlation functions (CCFs) to estimate correlated binding

Like the 2D ACF, the 2D cross correlation function (CCF) between either sfGFP-Zld or GAF-sfGFP and His2Av-RFP was computed through a fast Fourier transform protocol in Matlab. This 2D CCF was then used to fit a 2D model of cross correlation^42^:

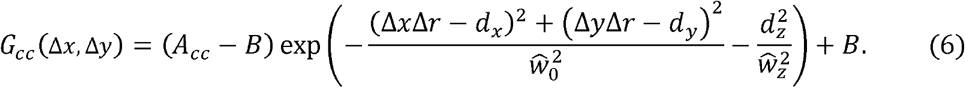

Because cross correlation uses two lasers, the two PSFs are different in size and their centers are generally not concurrent. In Equation 6, the parameters *d*_*x*_,*d*_*y*_,*d*_*z*_ represent the displacements between the centers of the two PSFs in the *x, y*, and *z* directions, respectively. The two displacements in the xy plane are adjustable parameters that can be estimated from the 2D CCF. The axial displacement was estimated by imaging fluorescent beads (see above), and our measurements resulted in the factor 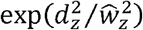 being roughly equal to 1.02, implying the factor could be safely ignored. As with the ACF, for robustness of fit, a small, adjustable background constant, *B*, was added. Fitting Equation 6 to the CCF data resulted in an estimate of the CCF amplitude, which is defined as:

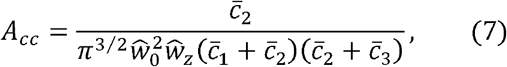

where the average PSF sizes are defined as 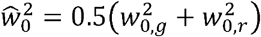and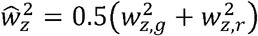, and the subscripts *g* and *r* denote the green and red channels, respectively, and where 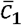 is the average concentration of the sfGFP-containing species that does not cross-correlate with the RFP-containing species (in the rest of the paper, this is the free pool plus the inactive pool); 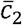 is the average concentration of the active pool, and 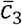 is the average concentration of the RFP-containing species that does not cross-correlate with the sfGFP-containing species, which include DNA-bound molecules, but could also include freely diffusing molecules.

To obtain *ψ*, the fraction of sfGFP-Zld or GAF-sfGFP correlated to His2Av-RFP, we calculate the ACF amplitude of the red channel, *G*_*red*_(Δ*x*, Δ*y*), then use it to fit Equation 3, yielding the amplitude, 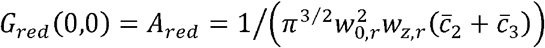. Then the ratio of *A*_*cc*_ to *A*_*red*_ gives:

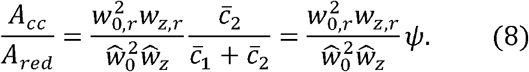

Therefore, we can calculate *ψ* as:

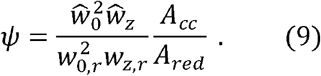

### Thermodynamic equilibrium model

When the binding of a TF to DNA is in equilibrium, the probability of the TF binding can be modeled as^84^:

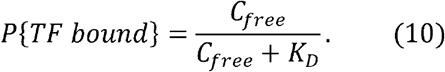

where C_free_ is the nuclear concentration of the free TF, K_D_ is the dissociation constant, *P{TF bound} = C*_*bound*_*/C*_*B*_,*C*_*bound*_ is the concentration of the bound TF, and *C*_*B*_ is the concentration of binding sites.

The equation was solved and adjustable parameters (dissociation constant and binding site concentrations) were determined using the least-squares fitting algorithm, lsqcurvefit, and the global optimization solver, MultiStart in Matlab.

To express the concentration of active or inactive bound TF as a function of total nuclear concentration, we used the conservation relation between different pools of the TF:

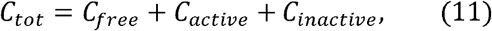

where C_tot_ is the total nuclear concentration, C_active_ is the concentration of the active pool, and C_inactive_ is the concentration of the inactive pool.

Using Equation (10) this can be written as:

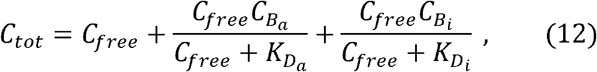

where 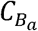 and 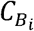 are the concentrations of binding sites for the active and inactive pool respectively. 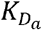 and 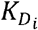 are the dissociation constants for the active and inactive pool respectively.

In the case of Zld, 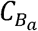 can be expressed as a function of C_free_ (see Figure S1D). Empirically, we modeled this relationship as a power law:

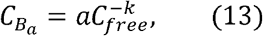

where *a* = 420 and *k* = 0.7 were the best-fit parameters (see Figure S1D). The value of 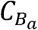 from Equation (13) can be substituted into Equation (12). Equation (12) can then be used to determine the concentrations of free TF corresponding to a given total concentration to construct the dose/response maps in Figures 2E, 3E. For these calculations, nc 10 is ignored due to its short duration and because data points near the start of each nc are excluded due to equilibrium not being established during these time points.

### Zld accessible sites

Pooled data from nc 8, 13, and 14 identified 12,135 Zld ChIP-Seq peaks^54^. We assumed that, during the early ncs, the chromatin is fairly open and all the ChIP-Seq identified sites are accessible for binding, such that the number of Zld sites accessible for binding during nc 11 is also assumed to be 12,135. Given that the nuclear volume during nc 11 is roughly 435 *μm*^3^;^85^, the concentration of Zld sites accessible for binding, averaged over the whole nucleus, is roughly,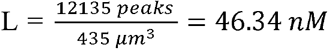. However, given that *Drosophila* is diploid, we set L = 2×46.34 = 92.68 nM. Using these estimates, Equation (10), and our data for nc 11, we arrive at K_D_ = 17 nM. Using this K_D_ and our data in Equation (10), the change in Zld sites accessible for binding over time was estimated.

### Residence time of Zld

Dissociation constants are typically defined as 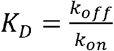; where k_on_ and k_off_ are the on and off rates for the TF-immobile structure interaction. For protein binding events in which the on-rate is limited by three dimensional diffusion, k_on_ is roughly 10^−4^-10^−5^nM^-1^s^-1^ (ref^86^). Using this rule of thumb for the k_on_ value, and *K*_*D*_ ∼ 20 *nM* (valid for both the active and inactive pools of Zld), the residence time, 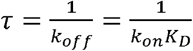 can be estimated to be: *τ* ∼ 8 *min* However, if the diffusivity is higher, as would be the case of one-dimensional sliding along the DNA^56^, k_on_ is closer to 10^−1^ nM^-1^s^-1^, resulting in *τ* ∼ 0.5 *s*.

The diffusivity of Zld obtained from our analysis, D ∼ 2 µm^2^/s. Using Smoluchowski equation^56^: *k*_*on*_ = 4 *πDba* where cross-section of the binding reaction, b = 0.34 nm and fraction of the. molecular surface of the protein that contains the reactive binding interface, a ∼ 0.2-0.5. Therefore, k_on_ ∼ 0.001-0.003 nM^-1^s^-1^, resulting in *τ* ∼ 19−49 *s*.

### Brightness analysis

The average molecular brightness, *Q*, of particles in the nucleus was calculated in the following manner. First, the average apparent brightness, *B*, of the sfGFP-containing particles was calculated as the ratio of the variance, *σ*^2^, of the intensity of pixels in the nuclei divided by their intensity,*I* :

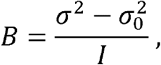

where 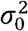 is the variance of a zero intensity image, which can also be found as the variance of the lowest intensity pixels in the image. Plots of *B* over nc 10-14 can be found in Figures S2A (Zld) and S4A (GAF).

Next, the *S*-factor, which relates apparent brightness and molecular brightness, was calculated as the slope of the best-fit line relating the microscope shot noise, 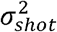, found in the pixels in the nuclei and the intensity of the nuclei^87^:

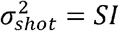

In practice, the shot noise is equal to the RICS ACF amplitude times the intensity squared, *AI*^2^, subtracted from the variance described above,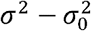. The shot noise and intensity at each time point in the time course was used to compute the best-fit line (see Figure S2B for Zld and S4B for GAF, for GAF). The equation used was 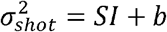, where *b* is a free parameter added for robustness of fit and was found generally to be close to zero. For Zld, *S* =363 and for GAF, *S* = 413.

Next, to convert the average apparent brightness into the average molecular brightness the following relationship was used^87^:

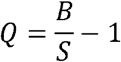

The plots of *Q* over nc 10-14 can be found in Figures S2C (Zld) and S4C (GAF).

Corrections for changes in brightness for GAF were performed in the following manner. First, a minimum value of *Q* was determined (*Q*_*min*_) and the brightness time course was normalized by Qmin: *q*(*t*) = *Q* _*min*_: *q* (*t*)/ *Q*_*min*_,with values of *q*(*t*) < 1being set to 1. Next, new values of the RICS ACF amplitude, *A*_*new*_, and of the fraction immobile,*ϕ* _*new*_, were calculated from *q* and the old values (*A* _*old*_ and *ϕ* _*old*_):

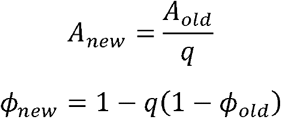

From these, corrected values of all concentrations, fraction free, and uncorrelated fraction were calculated. These corrections were performed at two levels. In the first level, we assumed all variations within nc 14 were within the range of being roughly constant. In other words, only variations outside of this range were corrected for, meaning that *Q*_*min*_ was set as the maximum value observed during nc 14 (see Figure S4C). This resulted in corrections made for the increase in brightness in the final point of nc 12 and the final four points of nc 13. While these corrections had a quantitative effect on our results, the corrected relationships between bound and free GAF are maintained (Figures S4D, E).

