## Supplemental information for "Bulk-level maps of pioneer factor binding dynamics during Drosophila maternal-to-zygotic transition"

Document S1. Figures S1–S5

Figure S1. Concentration of different pools of Zld, related to Figure 2

Figure S2: Molecular brightness of Zld.

Figure S3. Concentration of different pools of GAF, related to Figure 3

Figure S4. Molecular brightness of GAF.

Figure S5. Determination of axial displacement.

Video S1. Live imaging of sfGFP-Zld embryo from nc 10 to nc 14 used for RICS analysis, related to Figure 2. Acquisitions for the interphases of different ncs have been concatenated to generate the video, as mitoses were not used for the analysis.

Video S2. Live imaging of GAF-sfGFP embryo from nc 10 to nc 14 used for RICS analysis, related to Figure 3. Acquisitions for the interphases of different ncs have been concatenated to generate the video, as mitoses were not used for the analysis.

**Supplemental Figures**

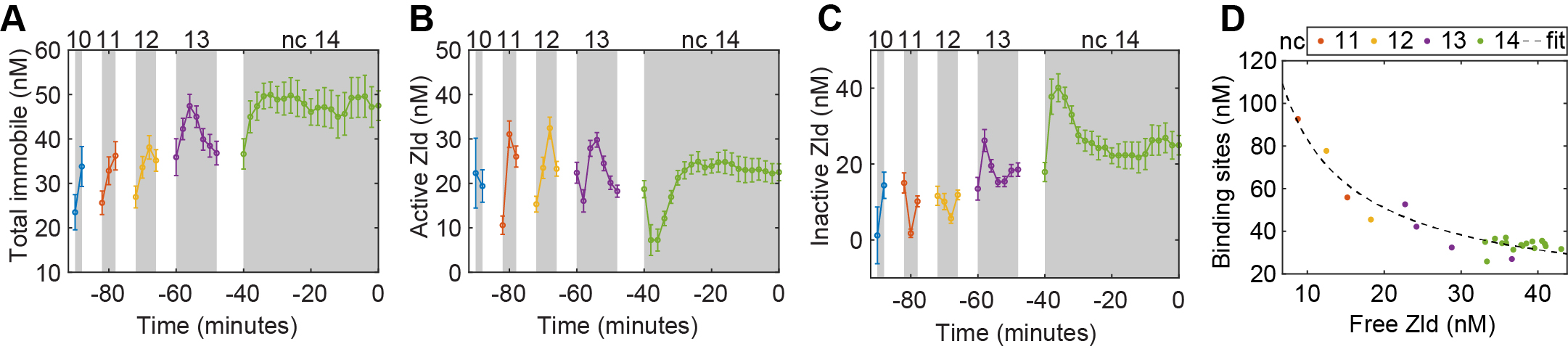

**Figure S1. Dynamics of concentration of different pools of Zld from nc 10 until gastrulation.**

(A) Immobile concentration.
(B) Active concentration.
(C) Inactive concentration. Line plots indicate the mean values for the measurements from 20 embryos. Error bars represent the standard errors of the mean.
(D) Power Law relation between concentration of binding sites and free Zld concentration.

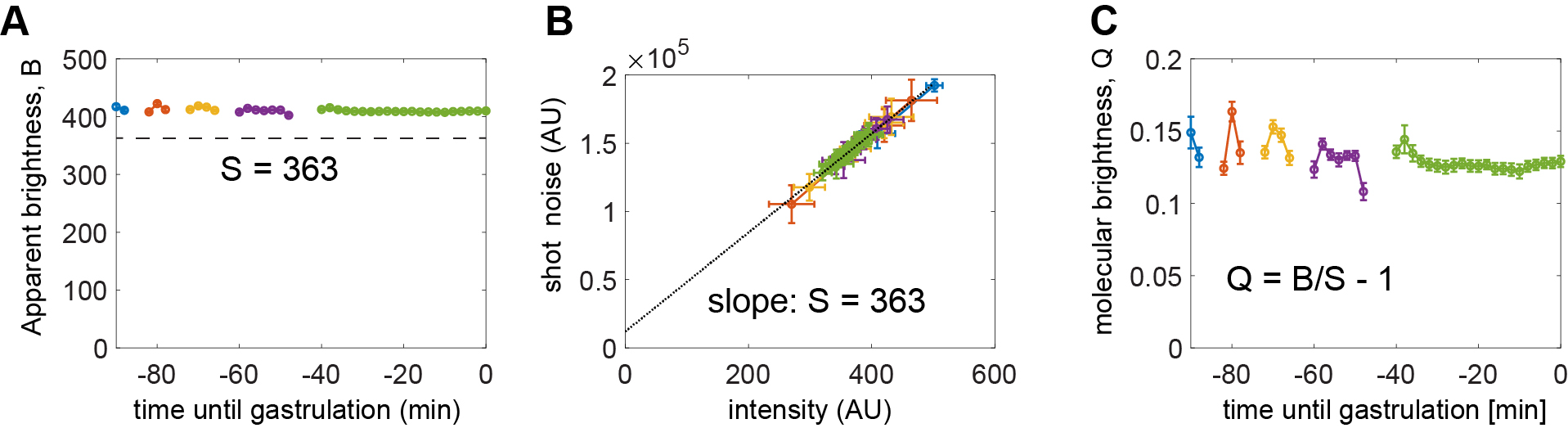

**Figure S2. Molecular brightness of Zld.**

(A) Apparent brightness, $B$, versus time. The horizontal line denotes the S-factor, calculated in (B)
(B) Shot noise versus intensity. The slope of this line is the S-factor that relates molecular brightness to apparent brightness.
(C) Average molecular brightness, $Q$, versus time. This brightness is roughly constant across all nuclear cycles with only a small variation. Error bars indicate SEM over all embryo time courses.

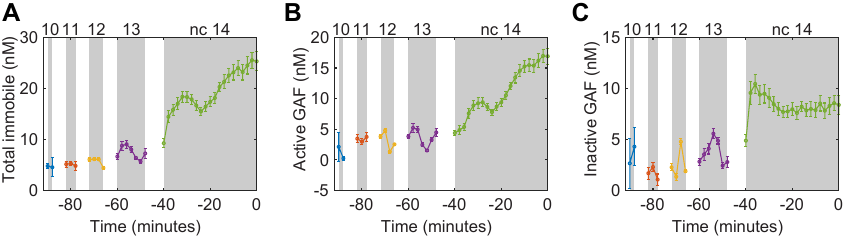

**Figure S3. Dynamics of concentration of different pools of GAF from nc 10 until gastrulation.**

(A) Immobile concentration.
(B) Active concentration.
(C) Inactive concentration. Line plots indicate the mean values for the measurements from 19 embryos. Error bars represent the standard errors of the mean.

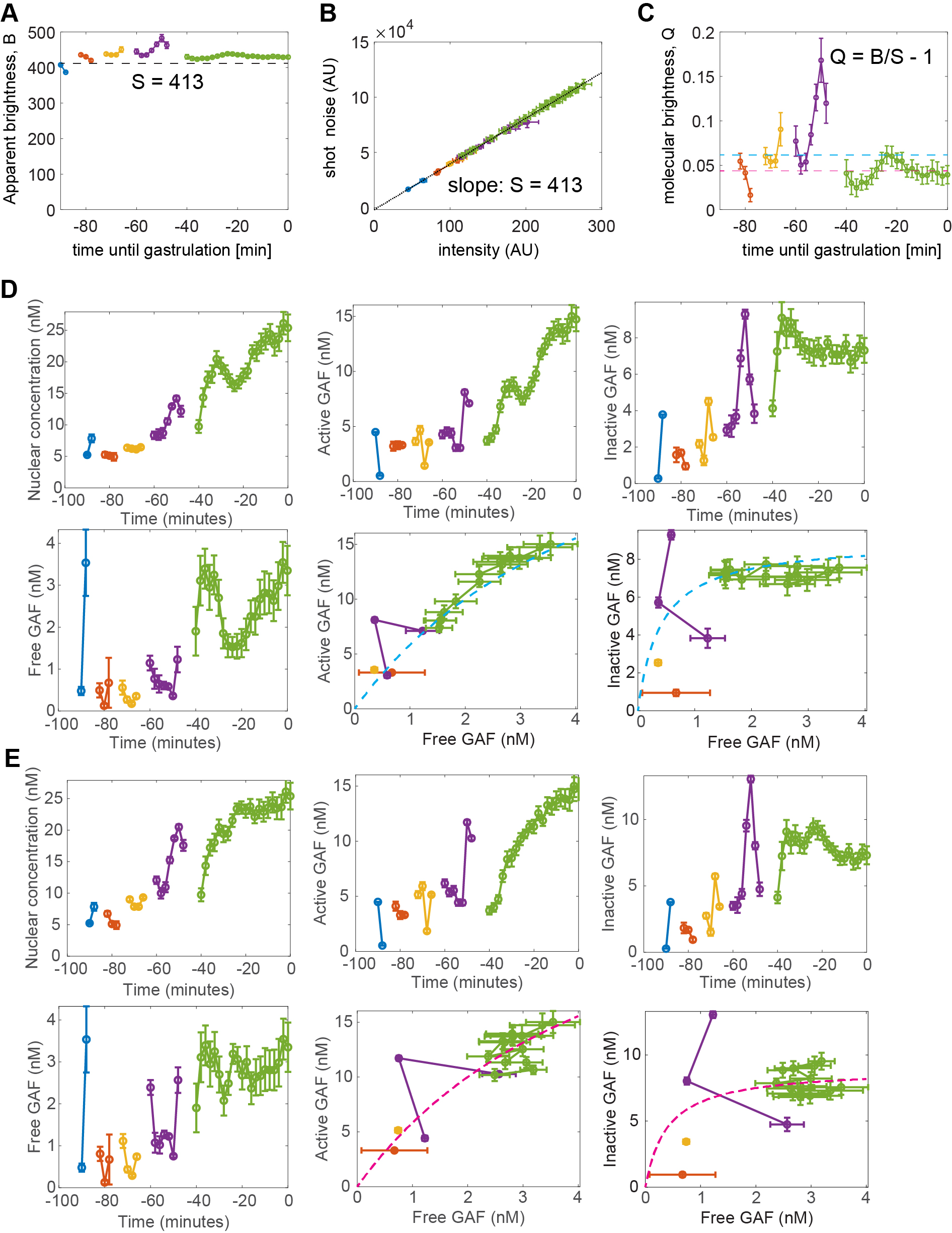

**Figure S4. Molecular brightness of GAF.**

(A) Apparent brightness, $B$, versus time. The horizontal line denotes the S-factor, calculated in (B)
(B) Shot noise versus intensity. The slope of this line is the S-factor that relates molecular brightness to apparent brightness.
(C) Average molecular brightness, $Q$, versus time. This brightness has some variation across the nuclear cycles, which may indicate corrections to concentration and immobile fraction are needed. Cyan horizontal line: corrections due to variations above this line are shown in (D). Magenta horizontal line: corrections due to variations above this line are shown in (E). Error bars indicate SEM over all embryo time courses.
(D) Corrections for dynamics of concentrations and the dose/response relationship due to variation above cyan line in (C)
(E) Corrections for dynamics of concentrations and the dose/response relationship due to variation above magenta line in (C)

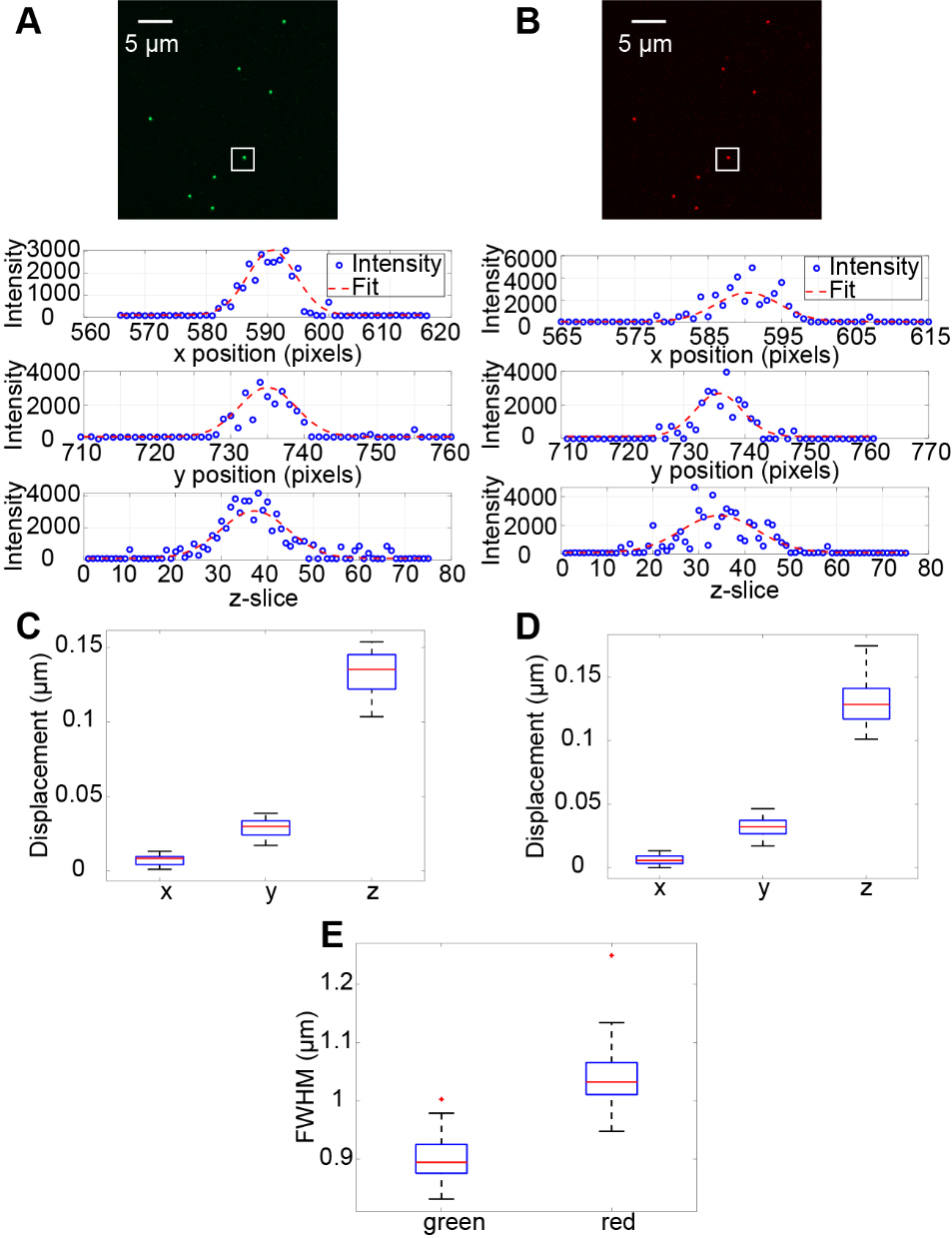

**Figure S5. Determination of axial displacement.**

(A) Representative image of beads in the green channel (top row). Gaussian fit to the intensity profile of the selected bead inside the white rectangle (in the top image) in the x, y, and z directions (bottom rows).
(B) Representative image of the same beads in the red channel (top row). Gaussian fit to the intensity profile of the selected bead inside the white rectangle (in the top image) in the x, y, and z directions (bottom rows).
(C) Axial displacement for the z-stack depicted in (A) and (B) (n = 8 beads).
(D) Axial displacement for multiple acquired z-stacks (n = 57 beads).
(E) FWHM of the beads in the two channels for multiple acquired z-stacks (n = 57 beads).
